# TIR-Learner v4: Accelerated annotation of terminal inverted repeat transposons

**DOI:** 10.64898/2026.07.21.739826

**Authors:** Kenji Gerhardt, Shujun Ou

## Abstract

The rapid expansion of high-quality, nearly complete eukaryotic genomes demands computational accelerations of existing bioinformatic infrastructure. TIR-Learner has been widely used for the *de novo* identification of Terminal Inverted Repeat (TIR) transposons, but suffers from slow runtime and a large memory footprint. Here we present TIR-Learner v4, a complete rewrite of TIR-Learner v3 and key supporting programs, which accelerates runtimes by two orders of magnitude and caps at a low RAM footprint irrespective of genome size. We demonstrate the scalability of TIR-Learner v4 by annotating TIRs in all 579 VGP Phase I genomes in around 4 hours.

## Introduction

The rapid growth of modern genomic data poses an ever-increasing challenge for bioinformatic software. In particular, the recent advances in long-read sequencing and assembly have made the production of highly accurate, highly complete eukaryotic genomes routine. In particular, the Vertebrate Genomes Project (VGP) aims to sequence all vertebrate species on Earth and has recently released its Phase I data freeze of chromosomal-level genome assemblies for 579 species^1^. Software methods that were previously effective in analyzing smaller genomes and lower volumes of data require improvement in their scalability.

Terminal inverted repeat (TIR) transposons are class II DNA-mediated transposable elements (TEs) prevalent in eukaryotic genomes. Annotation of TIR transposons has been limited by a lack of updates on existing software such as the GenericRepeatFinder (GRF) program^2^. Further highlighting the need for software updates is the active development of new software for the detection of TIR elements (in addition to other classes of TE), such as the HiTE^3^ program. TIR-Learner is a software pipeline for the *de novo* identification of TIR transposons that has undergone multiple iterations^4–6^ to improve performance and sensitivity. All versions share the same overall workflow: (1) a *de novo* structure-based search to find TIR candidates, (2) an optional homology-based classification of candidates found in (1), and (3) an ensemble classifier for fully *de novo* candidates. The initial repetitive element identification phase is implemented with the GRFMite subprogram in GRF. In the more recent v3 update^6^, the TIRvish subprogram in GenomeTools^7^ was added as an additional structural search to improve sensitivity. In all versions, the ensemble classifier includes random forest, K-nearest neighbors, AdaBoost, and convolutional neural network (CNN) sub-models, which all help remove false candidates. For brevity, we refer to these collectively as the “CNN module” of TIR-Learner.

While TIR-Learner is effective at identifying TIRs in principle, previous versions suffered from software design weaknesses that made the program scale poorly in both runtime and memory usage. These weaknesses in TIR-Learner v3 have prevented us from annotating TIR transposons within the VGP’s phase 1 release, including 579 nearly complete vertebrate genomes. TIR-Learner v3 was routinely unable to complete the annotation of large genomes such as axolotl (*Ambystoma mexicanum*, 28 Gbp) and African lungfish (*Protopterus annectens*, 40 Gbp). The theoretical strengths of TIR-Learner were therefore impractical to apply to these genomes, indicating a need for program redesign.

Here we present TIR-Learner v4, a vastly more computationally efficient reimplementation of TIR-Learner. TIR-Learner v4 is more than 10 times faster than v3 and operates consistently under 2 GB RAM per CPU thread. TIR-Learner v4 is capable of efficiently discovering TIRs in genomes of any size, which we demonstrate through its use to annotate TIRs in all VGP phase 1 genomes in just four hours.

## Results and Discussion

### Results

The overall program architecture of TIR-Learner v4 is a streamlined, modularized rewrite of the general logical workflow of TIR-Learner v3. Input genomes are divided into evenly sized work units of approximately 5 Mbp to facilitate work balancing across parallel threads (**Fig 1A**). Many logically similar tasks implemented in the v3 codebase, such as the retrieval of TIR candidate sequences from chromosomes and the post-processing loops of the TIRvish and GRFMite programs, were deployed as reusable modules that are shared across workflows.

**Figure 1:**
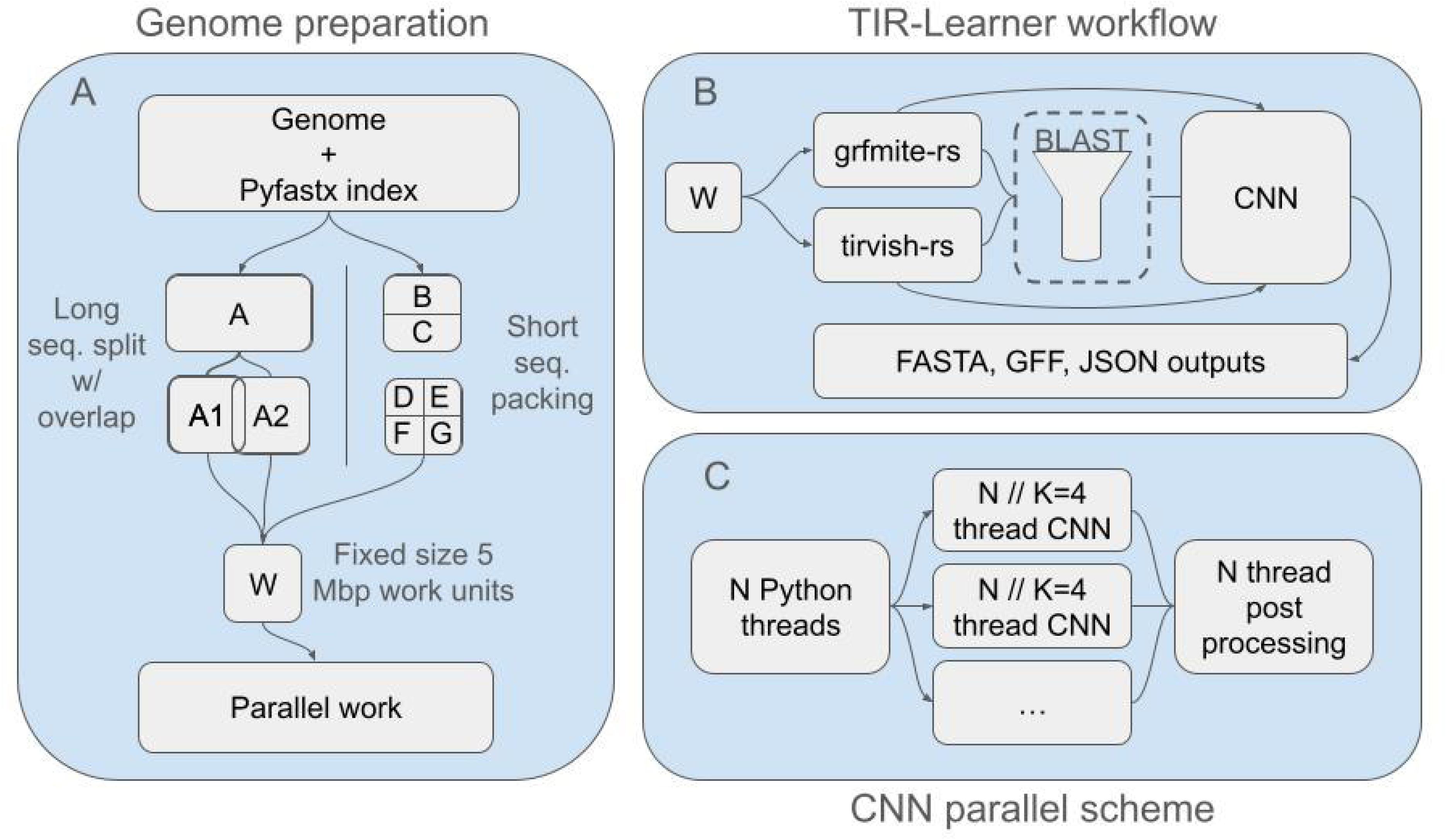
Program overview of TIR-Learner v4. (**A**) The genomeSplitter workflow. A genome index generated by Pyfastx is used to split long sequences into approximately evenly sized chunks no larger than 5 Mbp with 5,000 bp overlaps, while short sequences are packed into approximately evenly sized files also no larger than 5 Mbp. The genome is prepared as normalized work units (W), which prevent imbalances in workload size in subsequent processing. (**B**) TIR-Learner 4 modules and workflow. Work units are fed into grfmite-rs and tirvish-rs as subprocess calls. grfmite-rs and tirvish-rs are designed to handle parallelism internally, while filtering operations are managed in parallel via Python multiprocessing. TIR candidate sequences are optionally classified by BLAST ^8^ comparison to curated rice and maize TIR databases. Remaining (or all, if BLAST is skipped) candidates are filtered and classified into TIR families using TIR-Learner’s CNN module. Outputs are written in FASTA and GFF3 formats, alongside the JSON records of candidate identification and filtering. (**C**) The processing distribution scheme in the CNN step. Program threads are divided between Python processes and CNN backend OpenMP threads at a customizable ratio *K* with a default of 4, achieving a balance between efficient CNN performance and low RAM usage.

TIR-Learner v4 utilizes a more structured approach that isolates each relevant set of behaviors into separate modules that can be called independently and uses similar function signatures and parameterizations where possible. The outputs returned to TIR-Learner v4 from the tirvish-rs and grfmite-rs modules are unified into an identically structured JSON file, and the CNN module further outputs classification results to extend this JSON file (**Fig 1B**). Sequence retrieval and shared filtering components are likewise modularized and called rather than repeatedly implemented in each workflow. During the reimplementation, we also fixed several bugs identified in the v3 codebase and reported by users, including TIR identification and identity calculation (**Fig S1; Table S1**).

GRFMite is a subprogram of GRF that contributes the majority of program runtime in TIR-Learner v3. It contains algorithmic bottlenecks in multiple key processing stages that hinder sequence throughput. We reimplemented the C++ GRFMite in Rust as grfmite-rs. The majority of GRF behaviors are faithfully reproduced, but inefficient algorithmic choices in key bottlenecks have been replaced with more performant alternatives that achieve byte-identical outputs. Most significantly, the detection of initial seed regions of self-similarity and filtering of low-complexity sequences now utilize algorithms that avoid quadratic scaling, with time complexity reduced from O(n^2^) to O(n) (see methods). grfmite-rs is typically 16-32x faster in single-threaded comparisons compared to the original C++ GRFMite (**Fig S2**). Further, Rust’s rayon parallel architecture enables work-stealing that allows idling processors to “steal” workloads from other processors’ queues, which helps ensure that the TIR-Learner runtime stays roughly linear with genome size. This acceleration of GRF via grfmite-rs brings the overall program runtime of TIR-Learner from hours to minutes on most genomes and from days to a few hours for unusually large and/or repeat-dense genomes.

TIRvish is a subprogram of GenomeTools, which also contributes a substantial portion of program runtime in TIR-Learner v3. We reimplemented TIRvish into tirvish-rs using Rust and achieved substantial acceleration. The primary speedups are twofold: (1) TIRvish utilized a suffix array for the identification of target site duplications (TSD), which we replaced with a simpler brute force TSD search more appropriate for the small sequence comparisons; (2) TIRvish utilizes a forward dynamic programming algorithm for the calculation of TIR sequence similarity. This is an excellent choice but is less performant than more recent high-performance string edit distance calculation libraries like the Rust RapidFuzz^9^ library which is used in tirvish-rs. Collectively, we accelerate tirvish-rs to 6.09x faster on average (**Fig S3**) in single-threaded comparisons on a highly fragmented Pacific shrimp genome (contig N50 87.4 kb)^10^. The candidates identified by tirvish-rs are identical to the original TIRvish. Like grfmite-rs, tirvish-rs adopts the same Rust rayon parallel architecture with work-stealing to reduce tail effects and improve CPU utilization.

TIR-Learner v3 spawns multiple Python processes to distribute work in its CNN module. However, the Keras^11,12^ library used to implement each CNN search is also parallelized with OpenMP, which by default uses all available cores on a system. This is true for each separate process, leading to CPU overutilization. In TIR-Learner v4, the CNN module is accelerated by distributed, parallelized reading of TIR candidate sequences locally within each Python worker process and by controlling Keras OpenMP backend threading. This prevents processor contention and improves CNN processing performance by 1.4x to 5.9x (**Table S2**).

Compared to TIR-Learner v3, v4 achieves speedup ratios of 8.3x, 10.5x, and 12.5x with highly fragmented testing datasets of 312 Mbp, 640 Mbp, and the full/1.6 Gbp Pacific shrimp genome samples (**Fig 2A**; contig N50 87.4 kbp) at 48 cores. Since this genome was found to be pathological to the performance of both TIRvish and GRFMite, these speedups represented a significant achievement for acceleration in difficult genomic contexts.

**Figure 2:**
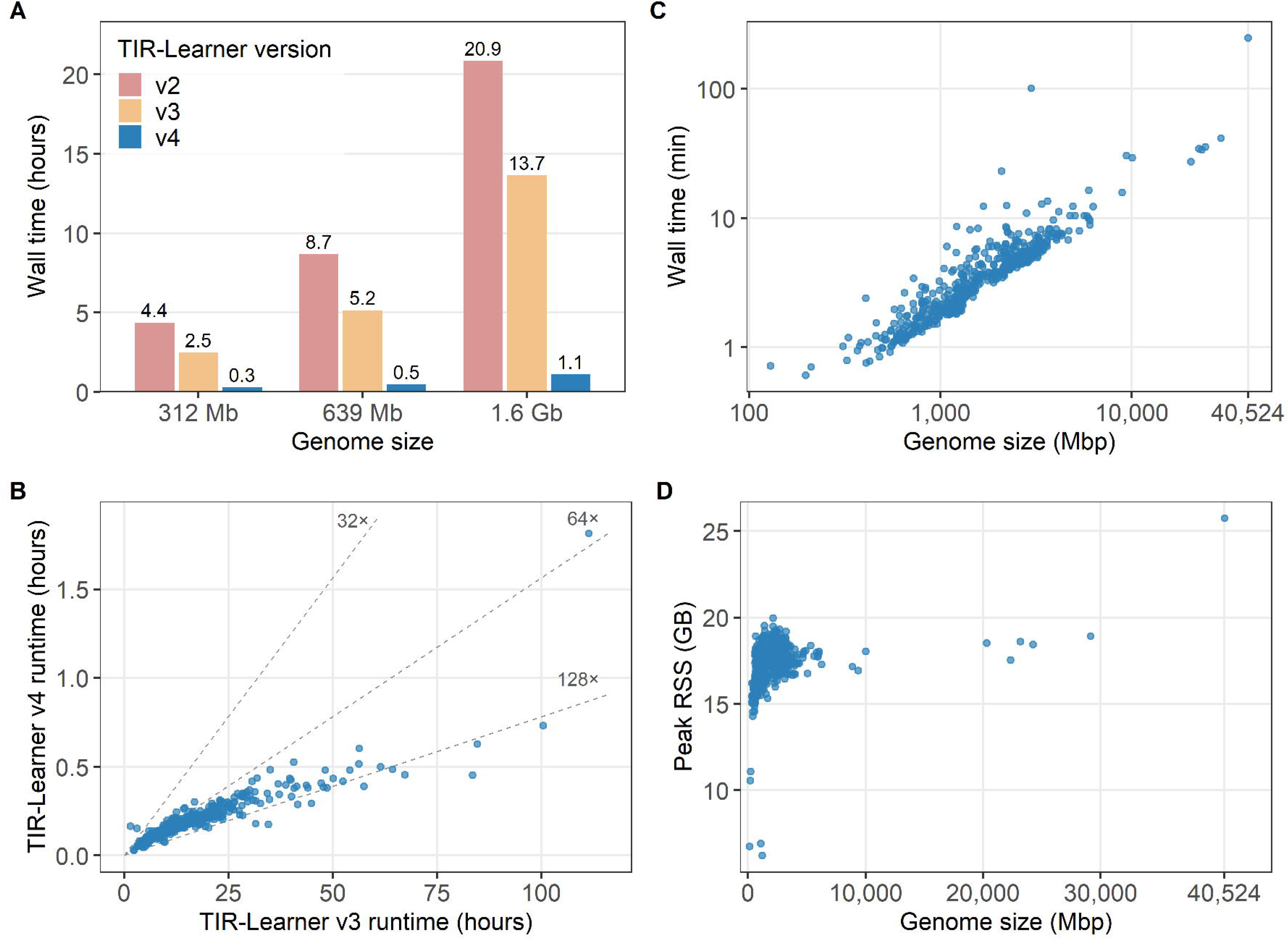
TIR-Learner v4 performance. (**A**) Comparison to previous versions of TIR-Learner on known pathological Pacific white shrimp genome fragments (contig N50 87.4 kbp). A 48-thread computing platform was used in this benchmark. (**B**) TIR-Learner v3 and v4 runtime comparison on 454 VGP genomes that v3 was able to process to completion. The dashed lines show the speedup ratio from v3 to v4. (**C**) Runtime and (**D**) peak RAM usage (Resident Set Size, RSS) of TIR-Learner v4 at 64 CPU cores on 579 VGP genomes. Each dot represents one genome. The X and Y-axes are log-scaled in **C**.

We benchmarked TIR-Learner v4 against v3 for runtime, CPU utilization, and RAM usage using the 579 VGP phase 1 genomes. Compared to TIR-Learner v3, v4 achieves speedup ratios of between 64x and 128x (95% CI 76.1 to 79.8x; **Fig 2B**; **Table S3**) on VGP genomes using matched CPU cores ranging from 8 to 48 (smaller genomes were allocated fewer cores for resource efficiency). Using 64 cores per genome on 24 concurrent jobs,TIR-Learner v4 completed within four hours and eight minutes for all 579 genomes (**Fig 2C**). Of the 579 genomes, 428 (74%) were completed in less than 5 minutes, and 550 (95%) were completed in less than 10 minutes. All but three genomes were completed in under an hour.

Runtime growth is approximately linear with genome size (**Fig 2C**). Variability in runtime beyond genome size is likely associated with the density of TIR candidates discoverable by grfmite-rs and tirvish-rs.

TIR-Learner v4’s fixed chunk size design results in a RAM usage plateau driven by the number of threads supplied to the program. Python parallel processes do not release memory until they complete; thus, each process holds the maximum amount of allocated RAM until process completion. The rapid growth of RAM between genomes <1 Gbp and ∼5 Gbp is the result of each parallel process trending towards a maximal but still reasonable allocation **(Fig 2D)**. Once every process has hit this limit (i.e., each process sees a TIR-rich genome chunk), RAM consumption plateaus at ∼0.33 GB RAM/thread.

## Discussion

In this study, we substantially improved the computational efficiency of TIR-Learner. The new program successfully annotated the 579 VGP genomes that harbor 1.22 trillion bases of sequence in only four hours of real time. This demonstrates that TIR-Learner v4 is a computational match for any forthcoming genome sequencing efforts. TIR-Learner v4 is capable of annotating thousands of complete genomes per day, and thus is no longer a computational impediment to TE annotation.

It is still challenging to classify TIR elements into superfamilies due to the lack of knowledge and sampling of true cases. Notably, the CNN classification model used by TIR-Learner v4 is the same model trained in TIR-Learner v2. This model was trained on curated TIRs from the Repbase. As VGP annotation efforts proceed, libraries of TEs, including TIRs that span essentially all vertebrates, will become available in non-model organisms. The forthcoming TIR and other TE libraries resulting from these efforts provide an opportunity to revise the core classification model at the heart of the program by retraining it with a much more comprehensive database of transposable elements in the near future.

## Conclusion

TIR-Learner v4 eliminates the computational roadblocks to the efficient annotation of TIR transposons in genomes. Compared to previous versions of the software, TIR-Learner v4 is faster, capable of processing genomes of essentially arbitrary sizes, and improves data transparency and reusability while maintaining the program’s overall workflow. TIR-Learner v4 is a substantial maturation of the program’s codebase, which will facilitate its use in the continuously growing and developing landscape of modern genomics.

## Code availability

TIR-Learner v4 is available at https://github.com/KGerhardt/TIR-Learner. genomeSplitter is available at https://github.com/KGerhardt/genomesplitter. grfmite-rs is available at https://github.com/KGerhardt/grfmite-rs. tirvish-rs is available at https://github.com/KGerhardt/tirvish_rs. grfmite-rs and tirvish-rs are also available on bioconda. All of these programs are licensed under GPL v3.

## Author Contributions

K.G. was responsible for code development, performance testing, and manuscript drafting. S.O. conceived the study, advised on project direction, and manuscript drafting and revision.

## Methods

### Work Balancing

TIR-Learner v3 uses a genome chunking scheme that divides a genome into 5 Mbp fragments, or one contig per fragment if the contig is shorter, to overcome limitations in the parallel efficiency of TIRvish and GRF. TIR-Learner v4 implements a more sophisticated genome splitting program, genomeSplitter (https://github.com/KGerhardt/genomeSplitter), which fragments the genome similarly with the additional feature of binning multiple short contigs into multiFASTA files containing approximately 5 Mbp in total. This prevents file count explosions in highly fragmented genomes and simultaneously improves parallel load balancing by ensuring there is no tail of small sequence fragments.

TIR-Learner v4 further embraces this divide-and-conquer scheme through the distribution of previously serial work to parallel workers. In v4, the TIR candidate subsequences identified by grfmite-rs and tirvish-rs are loaded locally from their respective chunks rather than their original sequences, and quality filtering is performed in this same local context instead of by the original sequences (e.g., chromosome). The CNN module further ensures workload balancing by divvying up particularly dense genome chunks into individual smaller batches, each with fewer than 20,000 TIR candidate sequences. Both intact and split CNN batches are passed to workers as small metadata dictionaries indicating which sequences the worker is responsible for, and the worker locally retrieves subsequences from the respective genome chunk files. This approach essentially eliminates data passing between the main process and the workers, enhances parallel efficiency, and naturally limits RAM used to only the candidates within individual genome fragments.

### GRFMite Rust reimplementation

The GRF program includes multiple command-line tools, and TIR-Learner only uses the Miniature Inverted repeat Transposable Element (MITE) detection module “GRFMite.”

The C++ GRF code uses multiple algorithms that are sensitive to highly self-similar, low-complexity DNA. Because essentially every sequence comparison in these cases matches many sequences, there is an exponential increase in the emission of candidate TIR sequences with corresponding exponential runtime increases. This worst case turned out to be common in the highly complete VGP genomes, with many containing complete assemblies of centromeres that tend to feature hundreds to thousands of centromeric repeats. These repetitive regions are often tiled in repeat units short enough to cause the C++ GRF code to produce and evaluate disproportionately large numbers of candidates in genome chunks containing these repeats. In TIR-Learner, this commonly resulted in a small handful of pathological genome fragments, each processing on a single thread, lingering long after all other non-pathological genome fragments had been processed with GRF.

The C++ implementation of GRF does contain multithreading as a command line option, but only about 50% of the program is actually parallel code. Consequently, it scales poorly with increasing thread counts and cannot exceed ∼2x faster runtimes regardless of the number of threads supplied. While supplying each GRF process with multiple threads would improve processing throughput for low-complexity sequences, it would substantially hurt performance in sequences without unusually heavy chunks.

As GRF was the primary rate-limiting component in TIR-Learner v4, we decided to improve this single GRF module. The GRFMite source code is about 200 lines, which made it an attractive target for selective improvement. We created grfmite-rs (https://github.com/KGerhardt/grfmite-rs), which is a Rust reimplementation of the C++ GRFMite program capable of producing byte-identical outputs. grfmite-rs uses faster but logically identical algorithms at essentially every computational stage, which is verifiable in the repo’s testing data using the C++ GRF backwards compatibility flags of grfmite-rs. The primary improvements in performance were alterations to how initial seed regions of self-similarity are identified and how low-complexity sequences are filtered. In the original C++ GRFMite, a two-pass approach is utilized in which self-similar regions are first identified as those with <2 mismatches across short windows which are then checked character by character in a second scan to produce a candidate list. grfmite-rs collapses these into a single pass where character comparisons are calculated from a two-bit encoded genome sequence via an Exclusive OR (XOR operation). In C++ GRFMite, sequence complexity is calculated (for the purpose of rejecting low-complexity sequences) with a loop including an O(n^2^) regex lookup step. grfmite-rs replaces this with an O(n) suffix automaton and adds an early exit condition where the calculation of sequence complexity stops as a sequence is guaranteed to be low complexity - an early exit condition that occurs soonest in the case of the most problematically self-similar bodies of sequence. Finally, the default output format of grfmite-rs is a compact, coordinate-only representation as opposed to the FASTA format of C++ GRFMite. grfmite-rs does support original FASTA format outputs for the purposes of ensuring identical program outputs between C++ GRFMite and grfmite-rs, but does not order outputs in the same way. Other program implementation changes are detailed in the GitHub repository.

### TIRvish Rust reimplementation

After the success of porting GRFMite from C++ to Rust, we decided to see if the TIRvish subprogram of GenomeTools (a library of mostly C scripts for a diverse range of genomics applications) was similarly amenable to acceleration in Rust. The TIRvish program was more tightly written from an algorithmic and implementation perspective. They made strong algorithmic choices in the most critical parts of the code. However, TIRvish still suffers from performance issues in a few places where solutions used by TIRvish were developed earlier in the lifetime of the GenomeTools ecosystem and were intended to serve the specific contextual needs of the programs they were originally designed for, such as LTR detection.

The TIRvish implementation consists of a five-stage filtering pipeline. In the first stage, candidates are identified via initial maximal exact matches (MEM) of at least 20 bp (default) and separated by no more than 5,000 bp (default). Second, initial candidates are fed through an implementation of the xdrop algorithm to identify full TIR candidate regions from those MEM seeds. This localizes regions as TIR “arms.” Third, the arms are checked for flanking target site duplication sequences of 2-13 characters, which must be exact matches and immediately follow the TIR arms of a candidate. Fourth, the TIR arm sequences are aligned to ensure that their total similarity is above a (default) cutoff of 80% identity. Finally, the surviving TIR elements are sorted and overlaps are resolved.

The primary performance weaknesses with the code as written were the choices of algorithm for TSD identification and TIR arm similarity checking. Both of these were inherited from GenomeTools’ LTR module. The TSD checking stage utilizes a locally constructed suffix array to find repetitive sequences. In the LTR context, where repeat units may be long and occur multiple times, this is an approach well suited to that specific problem space; however, TIRvish parameterizes TSDs as exact matches between 2 and 13 characters in length. For string comparisons of this length, the construction of a suffix array (especially one which will be used only once) is unnecessarily complex. The TIR arm comparison code in GenomeTools is an implementation of Levenshtein edit distance using a greedy diagonal front approach. This is a strong algorithmic choice for highly similar sequences, but is less performant when divergence increases (i.e., when a TIR candidate fails).

We developed tirvish-rs (https://github.com/KGerhardt/tirvish_rs) as a means of reducing the runtime of TIRvish. We first switched from a string representation of the genome sequence to a two-bit encoded representation to enable more rapid character comparisons in less space. We then implemented a brute force TSD search algorithm using this two-bit representation with bitwise XOR operations to rapidly detect sequence homologies. While the brute force approach would be less effective for very long sequences, it is a perfect fit for the 2-13 character space.

Second, we switched the calculation of Levenshtein distances to the RapidFuzz Rust crate’s banded bit-parallel edit distance function. This accomplishes two things: (1) it aborts the calculation of edit distance the moment that a sequence irretrievably falls below the 80% sequence similarity cutoff (default), saving time by not calculating the full edit distance of TIR arms guaranteed to be removed by the similarity filter, and (2) it is incredibly fast as a result of many low-level optimizations. Other program implementation changes are detailed in the GitHub repository.

The tirvish-rs code is designed for use in the context of TIR-Learner and expects that it will be run on 5 Mbp genome fragments. Where GenomeTools’ TIRvish used on-disk suffix array and longest common prefix array files, tirvish-rs stores these arrays in memory for each sequence for more performant access at the cost of inflated RAM usage. This would cause an explosion of RAM usage if applied to an intact genome, but safely maintains < 2 GB RAM/thread and averages 0.3 GB RAM per 5 Mbp fragment in the Pacific white shrimp testing data. The in-memory array in tirvish-rs also requires minimal additional working space to process even highly repeat-dense genomic fragments. To avoid memory inflation in use cases outside TIR-Learner v4, genomeSplitter should be used to preprocess the genome into 5 Mbp chunks before running tirvish-rs.

### TIR percent identity calculation bug fix

One of the few result-altering code changes in TIR-Learner v4 is a bug fix on how the program calculates TIR similarity as part of a filter between the tirvish-rs/grfmite-rs candidate-finding step and the CNN classification step. TIR-Learner v3 uses an incorrect implementation of Hamming distance - the edit distance between two strings of identical length. The implementation is incorrect because it calculates the edit distance character-by-character between the 3’ TIR and the reverse-complemented 5’ TIR. The v3 code exits this calculation as soon as similarity >80% and length >=10 characters and reports edit distance over only that span rather than over the full TIR. Usually, this is exactly 10 characters for a passing TIR pair, resulting in an artificial pileup of TIR identities at 100%, 90%, 80%, and with a few between 80% and 90%. Hamming distance is also inapplicable to strings of differing lengths, which happens when two TIR arms have an InDel. The v3 code dismisses this issue by terminating when the shorter arm’s length has been traversed, also producing an incorrect edit distance.

TIR-Learner v4 implements an improved TIR percent identity calculation step using the wavefront alignment algorithm from WFA2-lib^13,14^. Comparisons are always performed over full-length TIR candidate arms, and the switch to an alignment-based percent identity enables accurate comparisons in the presence of InDels. Since GRFMite and grfmite-rs only emit same-length TIR arms, the Hamming distance would always be calculable over these arms. However, TIRvish and tirvish-rs impose no such limitation and indeed about 75% of TIR candidates emitted by TIRvish contain TIR arms that differ in length by at least one base pair.

As a result, the majority of TIR candidates identified by TIRvish/tirvish-rs are incorrectly rejected by TIR-Learner v3. The TIR identity calculator in v3 thus systematically miscalculates TIR identity.

We quantified this effect by comparing the reported percent identity using TIR-Learner v3’s calculator to the new, wavefront-based TIR-Learner v4 calculator for TIR arms recovered in the human genome. As tirvish-rs produces a much smaller number of TIR candidate sequences (96k tirvish-rs candidates vs. 6.6 million grfmite-rs candidates in the human genome), the net effect appears small in the aggregate as only ∼1% of the total candidates are incorrectly rejected as a result of the bug. However, this fraction is also the substantial majority of tirvish-rs candidates. A total of 65,613 out of 96,476 tirvish-rs candidates from the human genome were incorrectly rejected according to the TIR-Learner v3 TIR similarity calculator. All of these rejected sequences are correctly retained in TIR-Learner v4.

Similarly, TSD identity calculation and filtering in TIR-Learner v4 is performed using the wavefront-based identity calculator.

### Other baseline algorithmic improvements

TIR-Learner v4 improves on essentially all other processing and filtering components of v3. Sequence and subsequence management by the BioPy^15^ SeqIO module in TIR-Learner v3 is replaced with the more performant Pyfastx^16^. The use of Pandas^17^ for data processing and Swifter^18^ for parallel operation in TIR-Learner v3 is eliminated in favor of logically equivalent but more performant NumPy^19^ operations and Python multiprocessing. Sequence composition assessment performed in v3 with calls to Python’s collections. The use of Python’s collections.Counter has been replaced with a NumPy array prefix sum implementation. This algorithm is enormously faster because it does not require loading subsequences at all, as sequence loci alone are sufficient after an initial precomputation of the array. Prefix sum filtering is particularly useful for grfmite-rs results, as the program produces highly overlapping, extremely redundant candidate sequence collections that can produce hundreds of gigabases from small genomes. Where the v3 approach separately processes each candidate sequence, the v4 approach requires only one pass over each ∼5 Mbp genome chunk and reuses the resulting array across all relevant candidates to obtain the same sequence composition information.

Finally, the Keras and PyTorch libraries that both TIR-Learner v3 and v4 use to implement their CNN classification steps are compiled natively with an OpenMP backend that, by default, utilizes all available threads on a system. This causes two major issues: first, it can sometimes access resources from more processors on an HPC deployment than are allocated to a job, causing cost overruns or interference on the node that reduces performance. Second, TIR-Learner already manages some parallelism in Python with its multiprocessing; each child process spawns a separate Keras invocation, each of which attempts to use all available threads. This leads to contention across every available core and dramatically reduces CNN classification speed. To avoid this, TIR-Learner v4 limits the processors available to each Keras invocation to a (default of) 4 while spawning 1/4th as many Python processes as the number of cores supplied to TIR-Learner v4. This utilizes all allocated threads without any contention during the expensive CNN stage while accepting modest efficiency losses for sequence loading and CNN post-processing. Full adoption of the Keras OMP backend was not selected as our testing showed quickly declining benefits with additional threads.

### Memory usage

TIR-Learner v3 adopted a strategy of mostly utilizing in-memory filtering operations compared to earlier versions of TIR-Learner that were reliant on more disk operations with increased program I/O. As a result, TIR-Learner v3 is about 30% faster than the previous TIR-Learner v2, even after accounting for its addition of the TIRVish program for TIR candidate identification, but has increased RAM usage. The development of TIR-Learner v3 used smaller genomes as its testing data, chiefly the highly fragmented Pacific white shrimp genome^10^ (1.6 Gbp in the assembly) and subsamples of this genome. At such a small scale, the sequence filtering approach is reasonably fast and has acceptable RAM usage; however, the all-memory approach scales poorly to larger genomes. As contig size increases (i.e., better quality and/or larger genomes), RAM use increases dramatically. This is further exacerbated by limitations in Python’s process-based parallelism that require duplicating data passed between the main and worker processes. The primary cause of the excessive RAM usage in v3 is the choice of a per-chromosome data blocking scheme. TIR candidate records are produced by both v3 and v4 on a per-chunk basis with no need for cross-fragment references aside from dereplication in overlap regions, yet TIR-Learner v3 aggregates these independent chunks into whole-chromosome collections for subsequence extraction and filtering, causing unnecessary RAM usage.

TIR-Learner v4’s fully local scheme avoids this limitation entirely: no substantive operations after the initial genome fragmentation step are performed at the whole-chromosome scale, only at the local 5 Mbp genome chunk scale. This scheme potentially saves hundreds of GB of RAM, and potentially saves hours of wall time for the serialization and deserialization step. Where data is passed between the main and worker processes, it is always in the form of small metadata summaries. Each worker’s total RAM requirement is thus reduced from TIR candidates over a whole chromosome (e.g., >2 Gbp for lungfish) to only those candidates from a ∼5 Mbp fragment. The switches from TIRvish to tirvish-rs and GRFMite to grfmite-rs are attended by higher RAM usage in each tool, especially in tirvish-rs. However, as these Rust reimplementations of the original program were developed for TIR-Learner v4, both are carefully designed to maintain a consistent RAM footprint per thread, resulting in an overall low RAM footprint.

### Output formatting and data governance

Finally, TIR-Learner v4 utilizes JSON-formatted summaries of tirvish-rs, grfmite-rs, and CNN processing outputs. Older versions of TIR-Learner discarded all information of candidate TIR sequences that failed to pass filtering criteria. TIR-Learner v4 records all candidate loci and filtering results, indicating clearly which candidates emerged from which program, which passed each filtering stage, and why pruned candidates were removed. TIR-Learner v4 also uses these summaries as part of a streamlined, compact checkpointing approach. An additional byproduct of the new approach is that temporary files are only produced on a per-worker basis as each genome chunk is processed; this, combined with the aggregation of short genome fragments into pooled chunks, can reduce the number of temporary files concurrently extant during TIR-Learners’ execution from potentially tens of thousands ((sum of long contig bases / 5 Mbp + each short contig) x 3 processing steps + working space per worker) to usually substantially under 10 thousand total (sum of genome bases / 5 Mbp x 1 processing step + working space per worker). This mostly eliminates the risk of file count limits on HPC deployments. After the TIR-Learner v4 execution, only 8 (or 12 in the homology mode) files remain as the final result. A summary of upgrades between TIR-Learner v3 and TIR-Learner v4 can be seen in Table S1.

### Benchmarking

To perform TIR-Learner v4 tests on the 579 VGP Phase 1 genomes, we used the Purdue ANVIL HPC platform. Each genome in the VGP collection was issued a 64-core allocation on an ANVIL standard memory node targeting a 64-core AMD Epyc “Milan” processor using an array job issuing up to 24 TIR-Learner instances concurrently. TIR-Learner v4 was executed using default parameters. We recorded wall time, CPU time, peak RSS, and CPU utilization for each job.

As TIR-Learner v3 is incapable of processing larger genomes due to RAM limitations, we compared the computational performance of TIR-Learner v3 and v4 over several fragmented Pacific shrimp genomes previously used for testing v3 and older versions. These runs were completed using 48 cores per job to match the testing approach used in the development of v2 and v3. The benchmark results of v2 and v3 were obtained from Ou et al. (2026)^6^.

We assessed the performance of grfmite-rs on an ANVIL node using 32 CPU cores. genomeSplitter was used with default TIR-Learner settings (5 Mbp chunk size, 5,000 bp overlap) to divide a European sprat fish genome into chunks before the performance test. Each chunk was processed sequentially by the C++ GRFMite and grfmite-rs on a single thread.

Parallelism was implemented per file via GNU parallel to prevent grfmite-rs from benefiting from its work-stealing mechanism. Program runtime was measured from invocation to termination on each chunk and for each GRFMite implementation. Outputs from grfmite-rs were compared to their respective C++ GRFMite results using diff; all were byte-identical.

We tested tirvish-rs with the same approach as grfmite-rs: 32 CPU cores on ANVIL with a pre-split genome and GNU parallel invocations of TIRvish and tirvish-rs, with each invocation receiving one thread. Instead of the European sprat genome, we used the full 1.6 Gbp Pacific white shrimp genome, likewise pre-split with genomeSplitter. As tirvish-rs does not support the GFF format of TIRvish, the contents of the outputs from TIRvish and tirvish-rs were parsed and compared using candidate coordinates. All candidates were shared with identical coordinate information between TIRvish and tirvish-rs across all genome chunks.

All VGP genomes used in this work were obtained via the VGP Phase 1 data freeze repository at https://github.com/VGP/vgp-phase1. The Pacific white shrimp genome was obtained at https://www.ncbi.nlm.nih.gov/datasets/genome/GCF_003789085.1/. The European sprat genome was obtained at https://www.ncbi.nlm.nih.gov/datasets/genome/GCF_963457725.1.

## Acknowledgements

This research was supported by the Ohio State University STEM Education Faculty Startup Award (S.O.) and the NSF ACCESS Discover computing grant (BIO250178 to S.O.). We thank the Purdue Rosen Center for Advanced Computing (RCAC) for providing technical support to the Anvil cluster.

## Supplemental Figures

**Figure S1:**
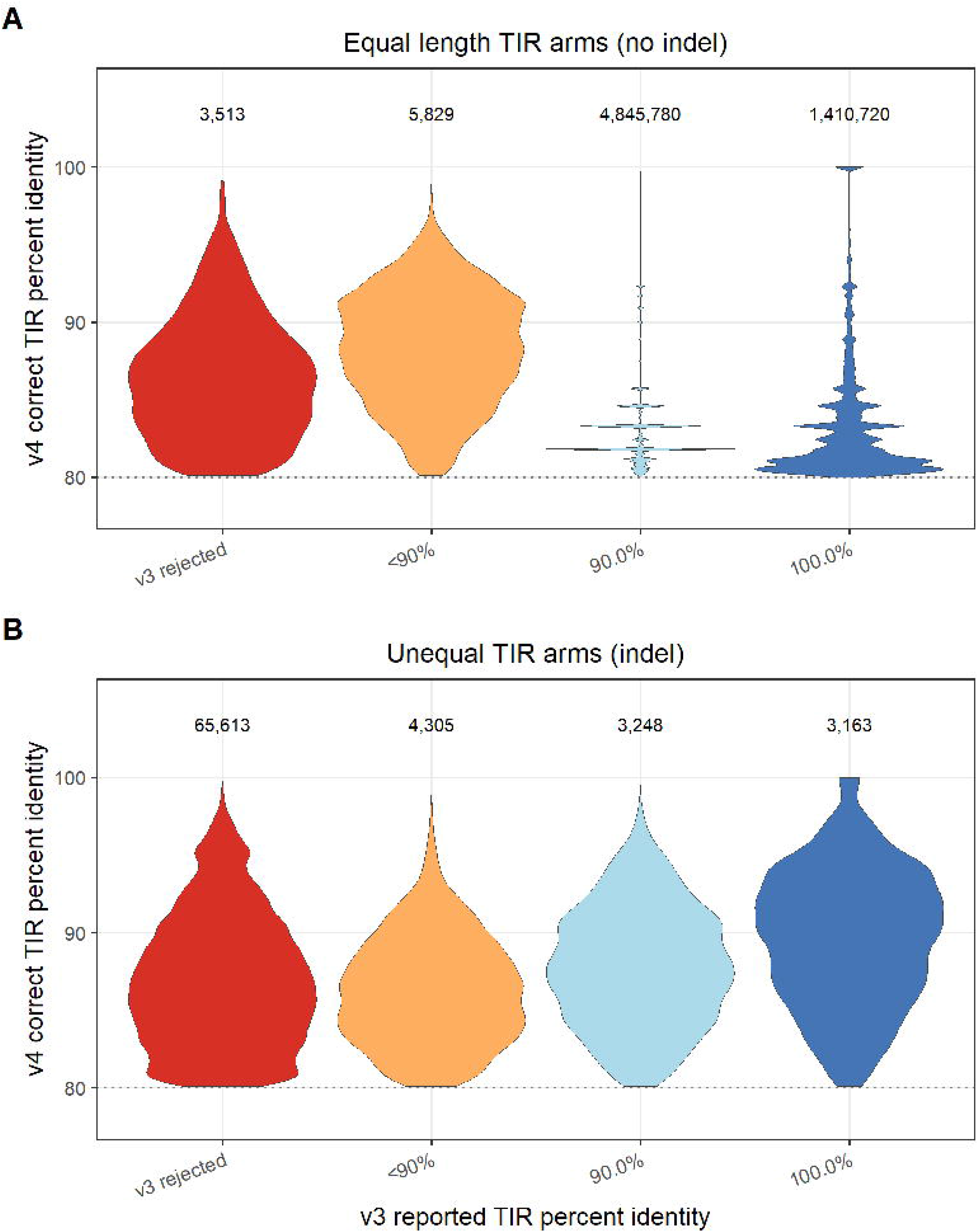
Result of TIR percent identity calculation bug fix between v3 and v4. **(A)** TIR-Learner v3 reported percent identity buckets vs. correctly calculated v4 percent identity on all human genome grfmite-rs and tirvish-rs TIR candidates without indels. Due to the identity calculation bug, TIR-Learner v3 reports a limited range of identity values (100%, 90%, and a few values approaching 80%). In contrast, TIR-Learner v4 reports continuous identity values. (**B**) v3 reported vs. v4 correct percent identity comparison for TIRs containing indels. Such cases are only produced by tirvish-rs and account for only ∼1% of total TIR candidates passed to CNN, but TIR-Learner v3 incorrectly rejects essentially all of these sequences.

**Figure S2:**
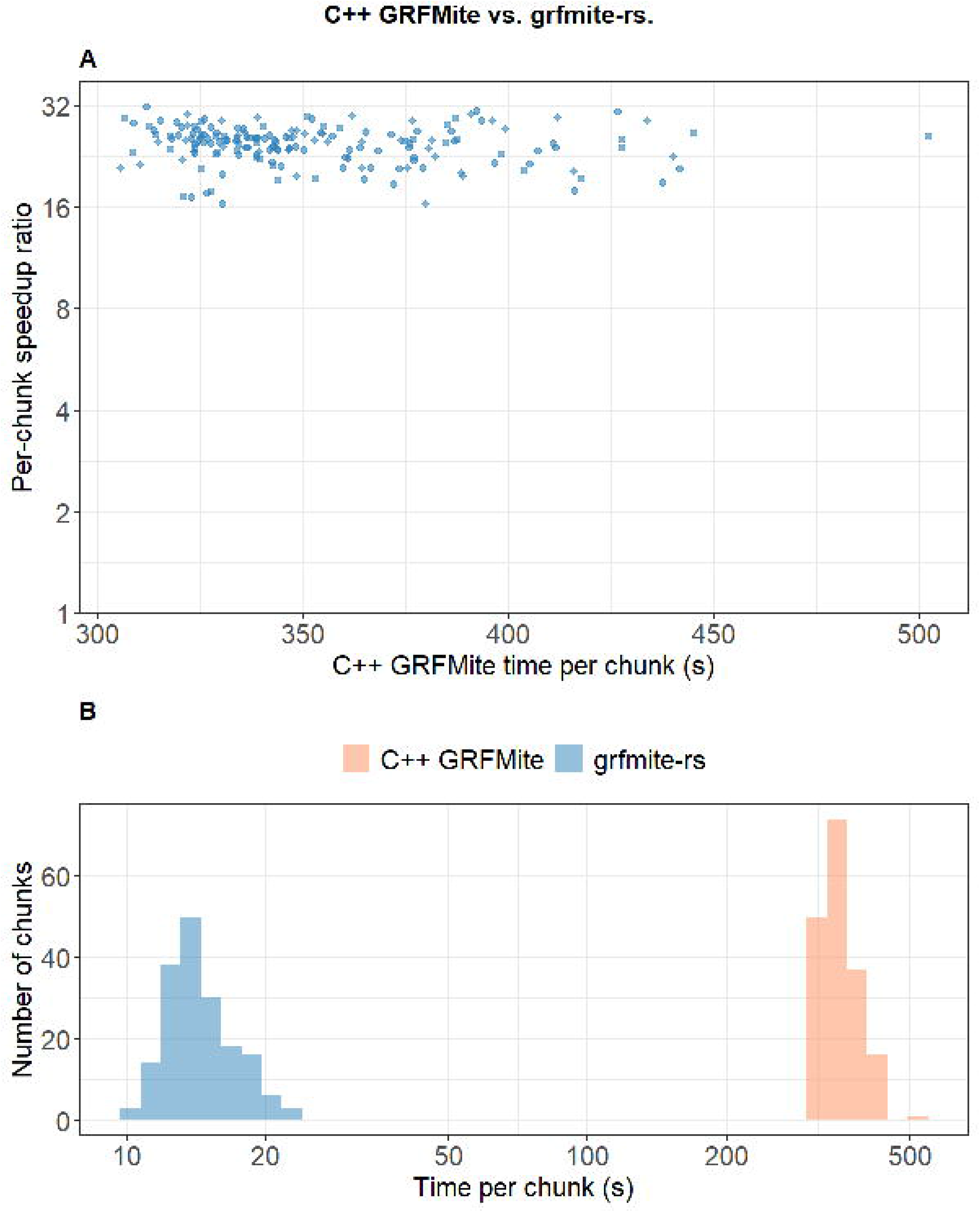
C++ GRFMite and grfmite-rs processing runtime comparison. (**A**) The C++ GRFMite runtimes of 5 Mbp genome chunks of the European sprat genome (<u>GCF_963457725.1</u>) (x-axis) and fold increase in speed for grfmite-rs on corresponding chunks (y-axis). Both programs were run with one processor per chunk. Challenging sequence chunks for C++ GRFMite are also challenging for grfmite-rs, but the acceleration is consistently 16x to 32x faster. The y-axis is scaled logarithmically. (**B**) Runtime distributions for the same genome chunks for grfmite-rs (blue) and C++ GRFMite (orange) in seconds. This x-axis is scaled logarithmically for readability.

**Figure S3:**
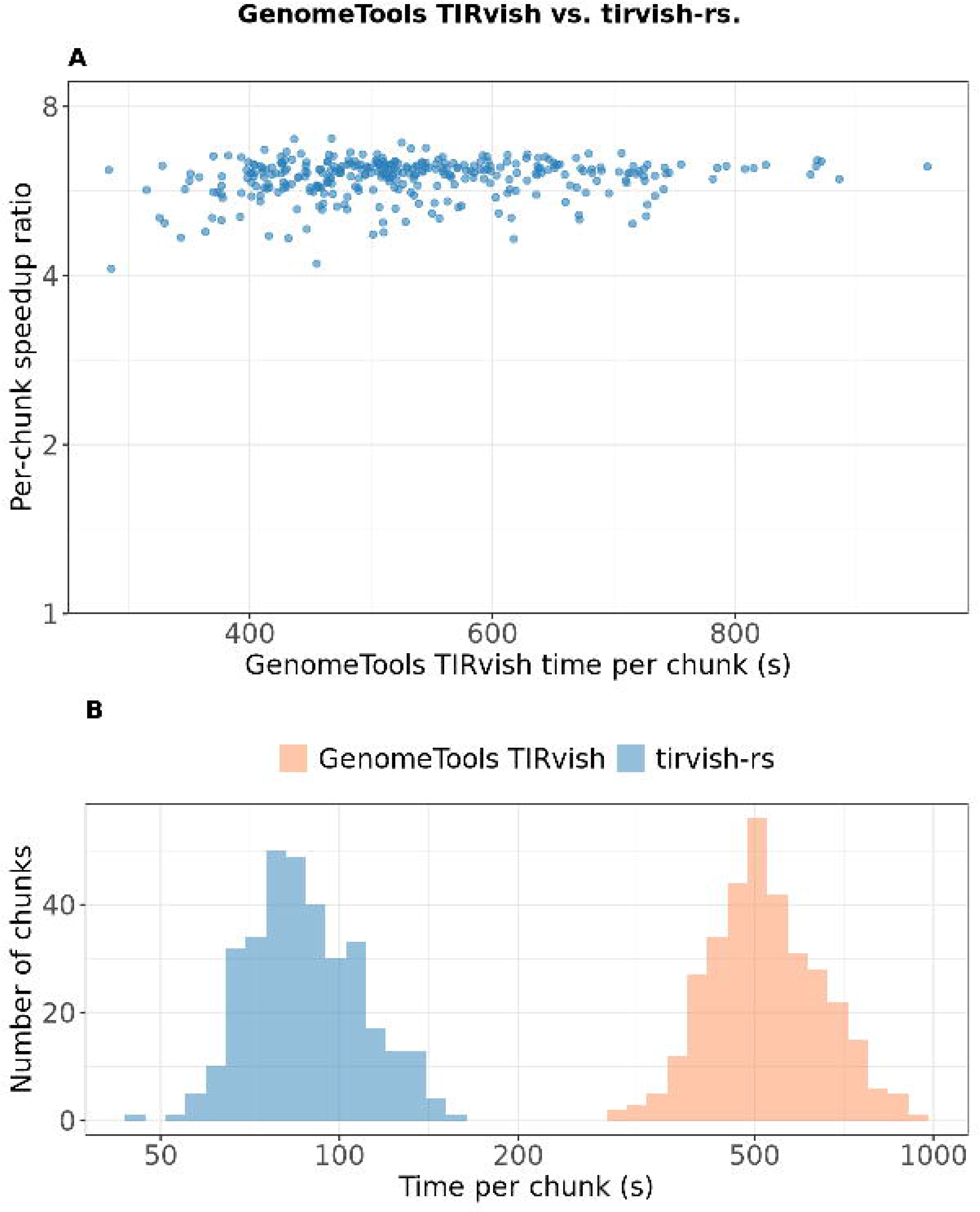
GenomeTools TIRvish and tirvish-rs processing runtime comparison. (**A**) GenomeTools TIRvish runtimes of 5 Mbp genome chunks of the 1.6 Gbp Pacific shrimp genome (GCF_003789085.1) (x-axis) and fold increase in speed for tirvish-rs on corresponding chunks (y-axis). Both programs were run with one processor per chunk. Acceleration is consistently 4x to 7x faster, stabilizing towards ∼6x overall as candidate density per chunk and runtime increase. The y-axis is scaled logarithmically. (**B**) Runtime distributions for the same genome chunks for tirvish-rs (blue) and GenomeTools TIRvish (orange) in seconds. This x-axis is scaled logarithmically for readability.

## Supplemental Tables

**Table S1:** Summary of weaknesses in the TIR-Learner v3 program and upgrades in TIR-Learner v4.

| TIR-Learner v3 | Deficit | TIR-Learner v4 | Advantage |
| --- | --- | --- | --- |
| Pandas + Swifter for data processing, parallelism | Data copying causes RAM issues, Pandas rows are an inefficient data structure for most TIR-Learner operations; Swifter parallelism is not well-suited to TIR-Learner use cases. | Numpy and bespoke multiprocessing parallelism, chunk-local design | Efficient data structures for each use-case; local design eliminates data copying, distributes I/O |
| BioPy SeqIO for genome index, sequence retrieval | Low-performance library | Pyfastx replaces all SeqIO instances | Highly performant; uses SeqTk script bindings |
| Residue counting per-subseq for sequence composition analyses | Redundant TIR candidate sequence pools result in hundreds of GB of unnecessary sequence processing per run. | Prefix sum arrays of AT, ambiguous base pair counts | One-pass calculation of arrays ~genome size bases processed per run, $O(1)$ cost to candidate-level counting |
| Uncontrolled CPU usage during Keras calls for CNN processing | Processor contention due to unmanaged conflict between Python process parallel and Keras OMP parallel | Controlled OMP parallelism with user-set ratio (default 4); load balancing 4 threads/Keras unit x N threads / 4 Python processes | No processor contention improves runtimes; distributed I/O overlaps read/write and processing |
| TSD and TIR filtering performed with brute force Python loops; early loop termination preventing correct %ID calculation | Nested Python loops produced quadratic scaling of runtime; incorrect identity values resulted from early termination. | Wavefront-based sequence alignment | TIR + TSD finding runtimes reduced to marginal program runtime, indel tolerance, and correctly calculated identity values. |
| Low optimization, parallelism within the GRFMite C++ | Parallelism implemented with OMP does not help | GRFMite rewritten in Rust with substantial algorithmic and | 16-32x faster runtimes on a single thread; work-stealing |
| implementation | the expensive serial components of the program that dominate cost in high-thread, workload-imbalanced cases. | parallelism improvements. | alleviates performance drop-off in highly repetitive regions. |
| GenomeTools' TIRvish uses contextually inefficient TSD finding and edit distance calculation approaches. | Reused TSD finding code optimized for LTR TEs was not a best-fit for TIR elements; Levenshtein distance calculation software has become more optimized since TIRvish was last updated | TIRvish was rewritten in Rust with algorithmic and parallelism improvements | 4-7x faster runtime on a single thread; work-stealing alleviates performance drop-off in highly repetitive regions. |

**Table S2:**
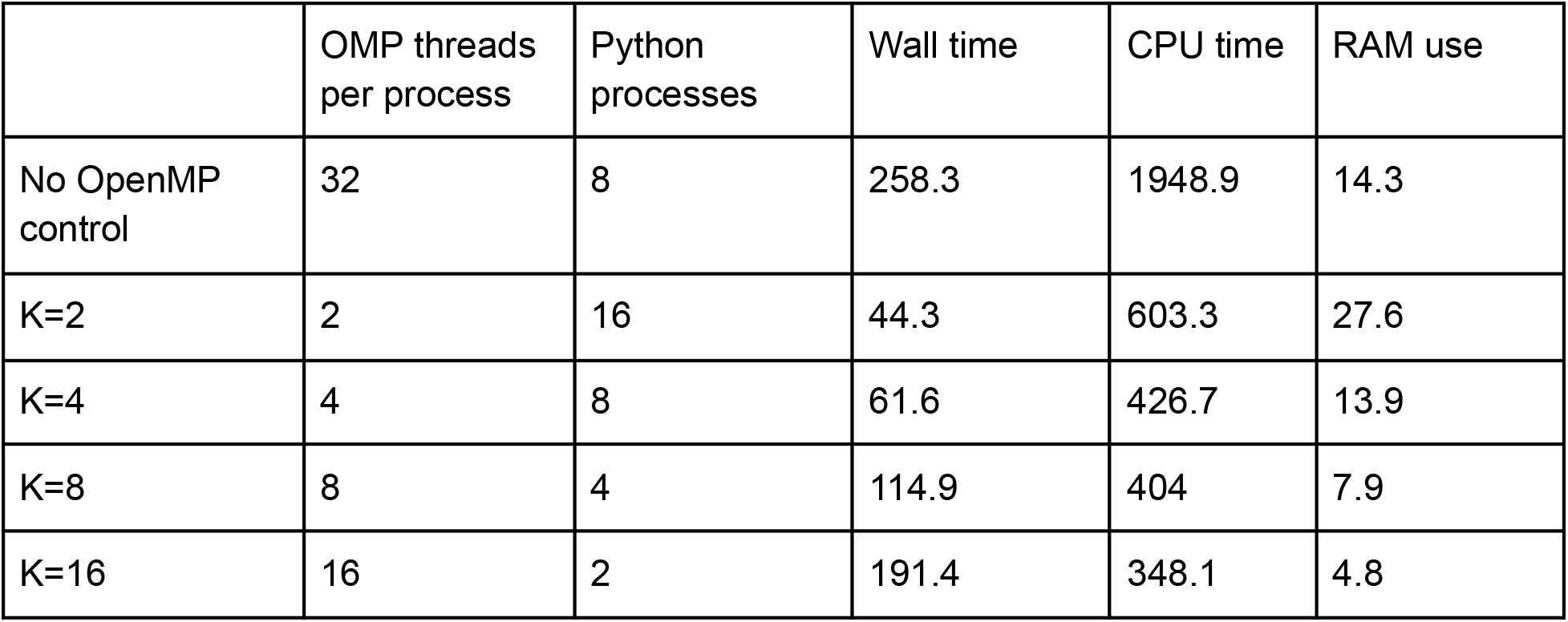
Effect of CNN backend threading control. Each job was run with 32 cores supplied on 498,934 TIR candidates; The K parameter determines the CNN cores per process.

## References

1. Formenti, G., et al. The Vertebrate Genomes Project Phase I: A global reference genome resource. bioRxivorg (2026) doi:10.64898/2026.06.24.732306.

2. Shi, J. & Liang, C. Generic Repeat Finder: A high-sensitivity tool for genome-wide DE Novo repeat detection. Plant Physiol. 180, 1803–1815 (2019).

3. Hu, K. et al. HiTE: a fast and accurate dynamic boundary adjustment approach for full-length transposable element detection and annotation. Nat. Commun. 15, 5573 (2024).

4. Su, W., Gu, X. & Peterson, T. TIR-learner, a new ensemble method for TIR transposable element annotation, provides evidence for abundant new transposable elements in the maize genome. Mol. Plant 12, 447–460 (2019).

5. Ou, S. et al. Benchmarking transposable element annotation methods for creation of a streamlined, comprehensive pipeline. Genome Biol. 20, 275 (2019).

6. Ou, S., et al. EDTA v2: enabling scalable TE annotation in animal genomes. bioRxiv 2026.07.01.735963 (2026) doi:10.64898/2026.07.01.735963.

7. Gremme, G., Steinbiss, S. & Kurtz, S. GenomeTools: a comprehensive software library for efficient processing of structured genome annotations. IEEE/ACM Trans. Comput. Biol. Bioinform. 10, 645–656 (2013).

8. Camacho, C. et al. BLAST+: architecture and applications. BMC Bioinformatics 10, 421 (2009).

9. Ye, A. et al. RapidFuzz: Accelerating fuzzing via Generative Adversarial Networks. Neurocomputing 460, 195–204 (2021).

10. Zhang, X. et al. Penaeid shrimp genome provides insights into benthic adaptation and frequent molting. Nat. Commun. 10, 356 (2019).

11. Chollet, F. Keras: Deep Learning for humans. https://keras.io (2021).

12. Ansel, J. et al. PyTorch 2: Faster machine learning through dynamic python bytecode transformation and graph compilation. in Proceedings of the 29th ACM International Conference on Architectural Support for Programming Languages and Operating Systems, Volume 2 929–947 (ACM, New York, NY, USA, 2024).

13. Marco-Sola, S. et al. Optimal gap-affine alignment in O(s) space. Bioinformatics 39, btad074 (2023).

14. Cleal, K., Kats, I., English, A., Lalli, J. PyWFA GitHub Repository. https://github.com/kcleal/pywfa (2022).

15. Cock, P. J. A. et al. Biopython: freely available Python tools for computational molecular biology and bioinformatics. Bioinformatics 25, 1422–1423 (2009).

16. Du, L. et al. Pyfastx: a robust Python package for fast random access to sequences from plain and gzipped FASTA/Q files. Brief. Bioinform. 22, bbaa368 (2021).

17. The pandas development team. Pandas-Dev/pandas: Pandas. (Zenodo, 2026). doi:10.5281/ZENODO.3509134.

18. Carpenter, J. Swifter GitHub repository. https://github.com/jmcarpenter2/swifterodo.3509134 (2018).

19. Harris, C. R. et al. Array programming with NumPy. Nature 585, 357–362 (2020).

